# Genomic divergence footprints in the bracovirus of *Cotesia sesamiae* identified by targeted re-sequencing approach

**DOI:** 10.1101/200782

**Authors:** Jérémy Gauthier, Philippe Gayral, Bruno Pierre Le Ru, Séverine Jancek, Stéphane Dupas, Laure Kaiser, Gabor Gyapay, Elisabeth A. Herniou

## Abstract

The African parasitoid wasp *Cotesia sesamiae* is structured in contrasted populations showing differences in host range and the recent discovery of a specialist related species, *C. typhae*, provide a good framework to study the mechanisms that link the parasitoid and their host range. To investigate the genomic bases of divergence between these populations, we used a targeted sequencing approach on 24 samples. We targeted a specific genomic region encoding the bracovirus, which is deeply involved in the interaction with the host. High sequencing coverage was obtained for all samples allowing the study of genetic variations between wasp populations and species. Combining population genetic estimations, the diversity (π), the relative differentiation (F_ST_) and the absolute differentiation (d_xy_), and branch-site dN/dS measures, we identified six divergent genes impacted by positive selection belonging to different gene families. These genes are potentially involved in host adaptation and in the specialization process. Fine scale analyses of the genetic variations also revealed deleterious mutations and large deletions on certain genes inducing pseudogenization and loss of function. These results highlight the crucial role of the bracovirus in the molecular interactions between the wasp and its hosts and in the evolutionary processes of specialization.

## Introduction

Interactions between organism of different species, such as between hosts and parasites, are thought to be one of the major evolutionary forces explaining lineage diversification and resulting into the current species diversity (Price 1980; Huyse *et al.* 2005). This diversification process has been particularly important in the Hymenoptera parasitoids for which about 50,000 species have been described (Quicke 2015). At the population level, the parasites evolve in a mosaic of interactions, theorized by the concept of the ‘geographic mosaic of co-evolution’ (Thompson 2005). This mosaic spreads over the host range, which comprise of all the permissive hosts allowing the parasite development, and which is variable both in space and time. Theories further suggest parasite populations should be locally more adapted to the hosts with which they most interact as reciprocal selection pressures should lead to arms-race (Ehrlich & Raven 1964) as symbolized by the Red Queen theory (Van Valen 1973). In this context different strategies exist depending on host range width. In the fields, we may find specialist populations interacting with a reduced number of host species (usually one) or generalist populations, also termed polyphagous or oligophagous, which can attack several host species. There are trade-offs associated with each strategy as specialists may be fitter on their particular host but dependant on its availability, while generalists may be more resilient to changes as they can exploit several hosts but are faced by more diverse resistance mechanisms. Under certain circumstances, the divergence of specialised host races may lead to speciation (Rice 1987; Via 2001; Peccoud *et al.* 2009).

Koinobiont parasitoid wasps are particularly dependant on their lepidopteran hosts because the larvae grow inside host caterpillars and from the outcome of the interaction only one of the partners will achieve its development and be able to reproduce (Gauthier *et al.* 2017). As they impact and limit host populations in the wild (Hawkins 1994), these wasps are used for biological control applications (Greathead 1986). This is the case of the generalist parasitoid wasp *Cotesia sesamiae* (Hymenoptera, Braconidae). Widely distributed in Africa, it parasitizes over twenty species from two lepidopteran families (Noctuidae and Crambidae), all belonging to the guild of cereal stem borers (Branca *et al.* 2012). But, previous studies revealed that this species presents a genetic structure comprising five major populations (Branca *et al.* 2017), which show variations in their host range (Branca *et al.* 2012) and in parasitic success on different hosts (Mochiah *et al.* 2002; Gitau *et al.* 2007). These 5 *C. sesamiae* populations include one population recently described as the new species *C. typhae* (Kaiser *et al.* 2015; 2017). All field samples from this lineage (N= 46 in Branca *et al.* 2017 and N= 35 in Kaiser *et al.* 2015) were strictly associated with larvae of the host *Sesamia nonagrioides* (Noctuidae), feeding on the host plant *Typha domingensis*. The five populations of *C. sesamiae* identified across Africa, show variations in geographic distribution, climate, symbiotic bacteria or host range. Among these factors, the main factor explaining genetic structuration is the host (Branca *et al.* 2017) revealing the strong selective pressure imposed by the hosts on the wasp populations.

*Cotesia* belong to the microgastroid group of braconid wasps in which the main virulence function is that evolved one of the most original parasitic strategies through the domestication of a virus (Webb & Strand 2005; Burke & Strand 2012; Herniou *et al.* 2013; Pradeu 2016; Gauthier *et al.* 2017). Bracoviruses derive from the stable integration of a large DNA virus of the family *Nudiviridae* in the genome of the wasps 100 Mya ago (Bézier *et al.* 2009; Thézé *et al.* 2011; Drezen *et al.* 2017). In the wasp genome, the bracovirus is composed of the proviral segments, which produce DNA circles carrying virulence genes. These segments are amplified, excised, circularized in DNA circles and packaged in bracovirus particles (Bézier *et al.* 2013). Female wasps inject bracovirus particles along with their eggs inside their hosts. The virulence genes encoded by the circles are expressed in the parasitized host caterpillar. They induce immunosuppression and other physiological modifications allowing the larval development of the wasp (Beckage *et al.* 1994; Labropoulou *et al.* 2008; Beckage 2012). In the genome of *C. sesamiae*, 17 out of the 26 proviral segments cluster at one locus, termed macrolocus, while the other segments are dispersed in the wasp genome. The segments encode 139 virulence genes grouped in 28 gene families (Jancek *et al.* 2013).

The adaptive role of the bracovirus was revealed with the study of two Kenyan populations of *C. sesamiae* (reviewed in Kaiser *et al.* 2017). Both populations vary in their capacity or failure to overcome the resistance of the host *B. fusca*. This capacity could be restored in the avirulent population with the transplantation of the calyx fluid, containing bracovirus particles, from the virulent population (Mochiah *et al.* 2002). Virulence against *B. fusca* was linked to allelic variation observed in the *CrV1* bracovirus gene (Dupas *et al.* 2008; Gitau et al. 2007; Branca *et al.* 2011). CrV1 was further found to harbour incongruent phylogenetic signal and contrasted signatures of positive selection from *histone-H4* and *ep2*, two other bracovirus genes when sampled in ecologically divergent wasp populations (Jancek *et al.* 2013). This suggests selection local adaptation of the wasps could be mediated by selection on different genes of their bracovirus. However, these studies on few genes give a restricted view of the potential implication of the bracovirus, which represent a collection of at least 139 virulence genes (Jancek *et al.* 2013). It is therefore necessary to investigate more widely the influence of the host range on the evolution of the bracovirus, which plays a central role in the parasitic success of the wasps. We developed a capture hybridization method (Hancock-Hanser *et al.* 2013) specifically targeting 300 kb of the genome of the wasp *C. sesamiae* containing the bracovirus. The targeted macrolocus as well as two dispersed loci were re-sequenced for 24 samples from *C. typhae* and different *C. sesamiae* populations of variable host range (Branca *et al.* 2017). We evaluated the efficiency and the robustness of the approach before undertaking phylogenomic analyses to assign the samples to the five previously identified populations (Branca *et al.* 2017). We then searched the molecular signature of adaptive evolution, by analysing differentiation between population F_ST_ and d_xy_ parameters and the ratio of substitution rates at non-synonymous and synonymous sites (dN/dS ratio), as well as pseudogenization. By tracking the evolution of the virulence genes encoded in the bracovirus of *C. sesamiae* at the population level, we revealed processes involved in host specialization and speciation. Overall our approach provides the first empirical insights on the genomic evolution of the domesticated bracovirus within the context of host parasite ecological interactions.

## Materials and Methods

### Biological material

Parasitized stem-borer host larvae were collected on wild plants in nine countries of Sub-Saharan Africa. For each lepidopteran larvae, GPS positions and altitude were recorded, as well as species identification and the host plant (Le Rü *et al.* 2006). Adult parasitoids were kept in absolute ethanol and identified to species based on genitalia morphology (Kimani-Njogu *et al.* 1997). Furthermore wasps from four parasitized caterpillars come from laboratory colonies originating from Kenya and maintained on the hosts on which they have been collected and regularly resampled for experimental purposes (Supplementary Table 1). Samples correspond to the pool of the adult *C. sesamiae* wasps emerged from these parasitized caterpillars. These pools, constituted of an average 30 gregarious *Cotesia* siblings, are not named “pools” to avoid confusions with the term used in Pool-Seq and composed of tens of individuals from the same population (Gautier *et al.* 2013). As the wasps are haplo-diploids, here, each sample represents at most the 3 genotypes of the female and male progenitors of the clutch studied. Pooling the clutch as one ‘sample’ was necessary to obtain the amount of DNA required for the targeted sequencing approach.

### DNA extraction, targeted enrichment and sequencing

Genomic DNA was extracted from the pool of wasp whole bodies using a Qiamp DNA extraction kit (Qiagen) with RNAase treatment following the manufacturer’s instructions and eluted in 200 μL of molecular grade H_2_O. DNA quantity was measured with a Qubit fluorometer (dsDNA BR, Invitrogen) and the quality was assessed by electrophoresis on a 1% agarose gel containing GelRed (Biotum). The DNA enrichment for regions of interest was performed using the SurSelect system (Agilent Technologies), for genomic DNA in solution (Mamanova *et al.* 2010). As several mismatches can be bridged during the hybridization step, this technology allows the capture of different sequences homologous to the reference.

The 300 kb targeted regions corresponded to 14 out of 26 *C. sesamiae* bracovirus segments of the Kitale population (Genbank accessions: HF562906-31), that were available at the time of the study design (Jancek *et al.* 2013; Bézier *et al.* 2013). Moreover, variations of the *histone-H4* and *ep2* genes (Genbank accessions: JX415828-44, JX430002-20) were integrated in the reference sequences to control for the effect of genetic variation on the capture efficiency. After enrichment, the samples were paired-end sequenced on an Illumina Hi-Seq 2000 instrument. All 100bp paired-end reads were sorted by samples using the MID tags and trimmed to remove low quality terminal bases and MID tags.

### Sequence analyses

Raw data analyses, including number of output reads and quality estimation, were done using the FASTX-Toolkit v0.0.13 (http://hannonlab.cshl.edu/fastx_toolkit/). Reads with more than 10% of low quality bases, i.e. below 20 Phred quality score, were removed and low quality bases were replaced by undetermined nucleotides, i.e. N. Reads of each samples were mapped to the Kitale reference sequence (accessions HF562906-31) using Bowtie2 v2.1.0 (Langmead & Salzberg 2012) with --no-mixed and --no-discordant options, which only allow paired-end reads mapping and thus avoid mismapping due to genomic duplications. Then, a pipeline including several steps of cleaning and filtering, based on recommendations from Kofler *et al.* (2011a) and Chen *et al.* (2012), was run to avoid biases and errors on allele frequency estimation. This pipeline first removed mapped reads with a mapping score lower than 20, using SAMTools v0.1.19 (Li *et al.* 2009), then locally realigned the reads using GATK IndelRealigner tool v3.4-0 (DePristo *et al.* 2011), and finally removed the PCR duplicates generated during the amplification steps of the target capture and sequencing using MarkDuplicates.jar from the Picard package v1.117 (http://picard.sourceforge.net). To evaluate and compare the efficiency of the targeted sequencing between the samples, the percentage of mapped reads, the base coverage, and the gene coverage, were measured with custom scripts including genomeCoverageBed.pl program implemented in the *bedtool* suite (Quinlan & Hall 2010).

The effectiveness and the completeness of the sequencing were estimated using a subsampling approach. For each sample, twelve random subsamplings of the mapped reads were done using DownsampleSam from the Picard package (http://picard.sourceforge.net) from 1% to 100% of reads. This process was replicated three times and at each condition, the diversity was measured to build a cumulative curve. To evaluate the sampling, we performed the same process by three random subsampling of samples in each population and by progressively increasing the number of sample from one to the maximum number. Global genetic diversity was estimated at each level using a custom script implementing Tajima’s π nucleotide diversity formula (Tajima 1983).

### Comparative genomics and phylogenomics

For each sample, a majority consensus sequence was generated using the “Generate consensus sequence” tool from Geneious suite v8.0.5 (Kearse *et al.* 2012) to extract the most common bases, i.e. most frequent alleles, with a minimum coverage of 20 reads. Then, these consensus sequences were aligned using Geneious aligner v8.0.5 (Kearse *et al.* 2012) and manually verified. The coding regions were extracted, concatenated, and a phylogenomic tree was built using the PhyML program (Guindon & Gascuel 2003) with the K80 substitution model determined using jModelTest (Posada 2008) and branch support was measured based on 1,000 bootstrap iterations.

### Population genetic analyses

For the analyses at the population level, a dataset for each population was created combining the data of the corresponding samples. In the absence of information on the emergence composition (i.e. ploidy level linked to the mating and emergence sex-ratio) the alignments from each sample were normalized by subsampling to a fixed coverage of 100X as recommended by the authors using subsample-synchronized.pl from the POPOOLATION2 pipeline v1.201 (Kofler *et al.* 2011b). Once normalized, the samples were merged by summing the coverage of each allele at each position. This allows to give each sample the same weight in the final population dataset. On these population datasets, different indexes were estimated to characterize genetic diversity and differentiation. All these estimations were performed at the level of bracovirus genes rather than on the entire bracovirus loci or using sliding windows, because transposable elements present in the intergenic regions of the bracovirus captured copies present elsewhere in the wasp genome, thus biasing the estimations. The nucleotide diversity of each gene was estimated using a custom script implementing Tajima’s π nucleotide diversity formula taking the gene length into account (Tajima 1983). The independence from gene size of these estimators was verified using correlation tests, which were all non-significant (Spearman’s rank order test, *p*-values > 0.06). The differentiation index (F_ST_) between populations was estimated with the POPOOLATION2 pipeline v1.201 (Kofler *et al.* 2011b) using the F_ST_ estimator developed by Weir and Cockerham (1984) allowing the comparison of populations of different sample size. For each estimation, the following filters were used: minimum base quality of 20, minimum coverage of 20, minor allele count of 1 and minimum covered fraction of 0.6. As within-population diversity can influence the measure of the relative differentiation (Cruickshank & Hahn 2014), the absolute differentiation (d_xy_) was measured in the population consensus sequences on each gene with the POPOOLATION pipeline v1.2.2 (Kofler *et al.* 2011a). For each index, Tukey’s box and whiskers plot method was applied to determine outlier genes, based on the global distribution of values rather than the last quantile approach. Outlier genes were thus defined as significantly different from the others when their index values were higher than the upper fence of the whisker (>Q3+1.5*IQ, with Q3 the third quartile and IQ the interquartile range). All statistical tests and graphics were done in R (R Core Team 2013). To investigate the genes under divergent selection, we used a phylogenetic method based on the ratio between synonymous and non-synonymous substitution rates (ω=dN/dS). The dN/dS ratios were estimated using the Branch-Sites REL model implemented in HyPhy testing all the branches (Pond *et al.* 2005). Significant estimations were confirmed using a second method, the Branch-Sites model (model = 2 and NSsites = 2), implemented in PAML (Yang 2007). Significance was evaluated by the comparison of a general model allowing positive selection (ω > 1) with a neutral model (ω fixed in 1) with a likelihood ratio test (LRT) based on the 2ΔlnL statistics (Yang 1998).

### Pseudogenization and gene loss

Bracovirus gene alignments were manually examined to identify insertions, deletions and non-sense mutations inducing premature stop codon. Only fixed mutations, shared by all samples of a given population, were recorded. Furthermore, the analysis of the gene coverage identified large deletions corresponding to regions without any reads. These deletions were confirmed by the assembly of the reads from each side of the deletion with Roche Newbler software v2.8 (Margulies *et al.* 2005) and the manual verification that the contigs overlapped the deletion gap.

## Results

### Efficiency of the targeted sequencing

The strategy consisting in the enrichment of the targeted bracovirus allowed the production of an average of 3.58 million paired-end reads per sample with a range of 2.89 to 5.14 million. On average, 43.66% of reads (range: 27.13%-49.90%) were mapped on the reference sequences after all filtering steps. Genetic distance did not affect sequence capture as 37.65% of reads from the outgroup species, *Cotesia flavipes*, could be mapped, which is well within the range and broadly similar to result from the reference sample *C. sesamiae* “Kitale” (43.89%; Supplementary Table 2).

The coverage of the targeted regions was above 441X, with an average of 649X for all samples including *C. flavipes* outgroup. Fine scale analysis of the coverage within the sequenced targeted segments revealed intergenic regions with extremely high coverage. Transposable elements (TE) were identified in the bracovirus segment and already reported in *C. congregata* species (Bézier *et al.* 2013). The excessive coverage presumably resulted from the targeted capture of TE copies present elsewhere in the whole wasp genome. As these additional TE reads could skew the genetic diversity of intergenic regions within the bracovirus loci, all further analyses excluded intergenic and TE sequences and were solely performed at the level of the 98 captured bracovirus genes..

To evaluate if the genetic diversity of reference sequences influenced DNA capture efficiency, two genes for which we had several genetic variants, respectively ten variants for the Histone H4 gene and thirty for the EP2-2 gene, were integrated in the targeted capture process, increasing the diversity of the sequence matrix for these genes. After capture and sequencing, both genes showed higher coverage than other targeted genes (Wilcoxon sum rank test p-value: 2.2^-16^ and 1.069^-07^; Supplementary Figure 1a), due to the sequence abundance of the capturing matrix; i.e. more abundant matrix in the capture mix, gives, in proportion, more captured sequences. However, in both cases increased coverage did not affect genetic diversity, which was not significantly different from other genes (Wilcoxon sum rank test p-value: > 0.05; Supplementary Figure 1b). This suggests that the matrix, based on a single sequence, is able to capture all the diversity present in the samples. Conversely, as neither Histone H4 nor EP2-2 showed lower diversity than other bracoviral genes, this implies that higher sequencing did not affect variant amplification, which could have artificially reduced the diversity captured. Therefore, coverage variability between genes can be attributed to random events during the capture rather than specific technical artefact.

To further test if the sequencing depth achieved in the experiments was sufficient to capture the genetic diversity present in each sample, we performed read subsampling (Supplementary Figure 2a). The cumulative curves showed that for all samples 10% to 30% of the reads were sufficient to reach a plateau of genetic diversity. Therefore adequate sequencing depth was achieved in the experiment. A similar approach was done at the population level to assess the representativeness of our samples, by increasing the number of samples for each population (Supplementary Figure 2b). For each population the curve reached a threshold of genetic diversity between three and four samples. This suggests that the genetic diversity contained in these samples is sufficient to characterise the population. Our sampling size is thus sufficient to represent *Cotesia* population diversity.

Altogether, we found on the one hand that the study design could capture the diversity present in homologous regions to the bracovirus target and on the other hand, that a complete absence of sequence data for a target region would likely indicate a genuine loss rather than a technical artefact.

### Phylogenetic analyses

To determine how the 24 *Cotesia* samples used for the targeted resequencing experiment relate to one another, we used the majority consensus sequence derived from the reads mapping on the bracovirus genes, as it reflects the main phylogenetic signal present in the sample. Five phylogenetic lineages could be clearly delimited (Figure 1b), *C. flavipes* the outgroup, the recently described species *C. typhae* and three *C. sesamiae* populations, in agreement to previous population genetic studies (Branca *et al.* 2017; Kaiser *et al.* 2015). Furthermore, the sample G4916 from the Endebess locality is genetically isolated and probably represents a single sample from a fifth population. However as we only had one sample from this fifth population, we could not carry out population-based analyses, as its diversity would not be representative. Geographic distribution appears to play a role in the genetic structuration of *C. sesamiae*. Populations 1, 2 and 3 are clearly distributed in different regions of Africa although with some overlap. *C. typhae* has also been reported in localities shared with *C. sesamiae* population 1 (Figure 1a).

**Figure 1.**
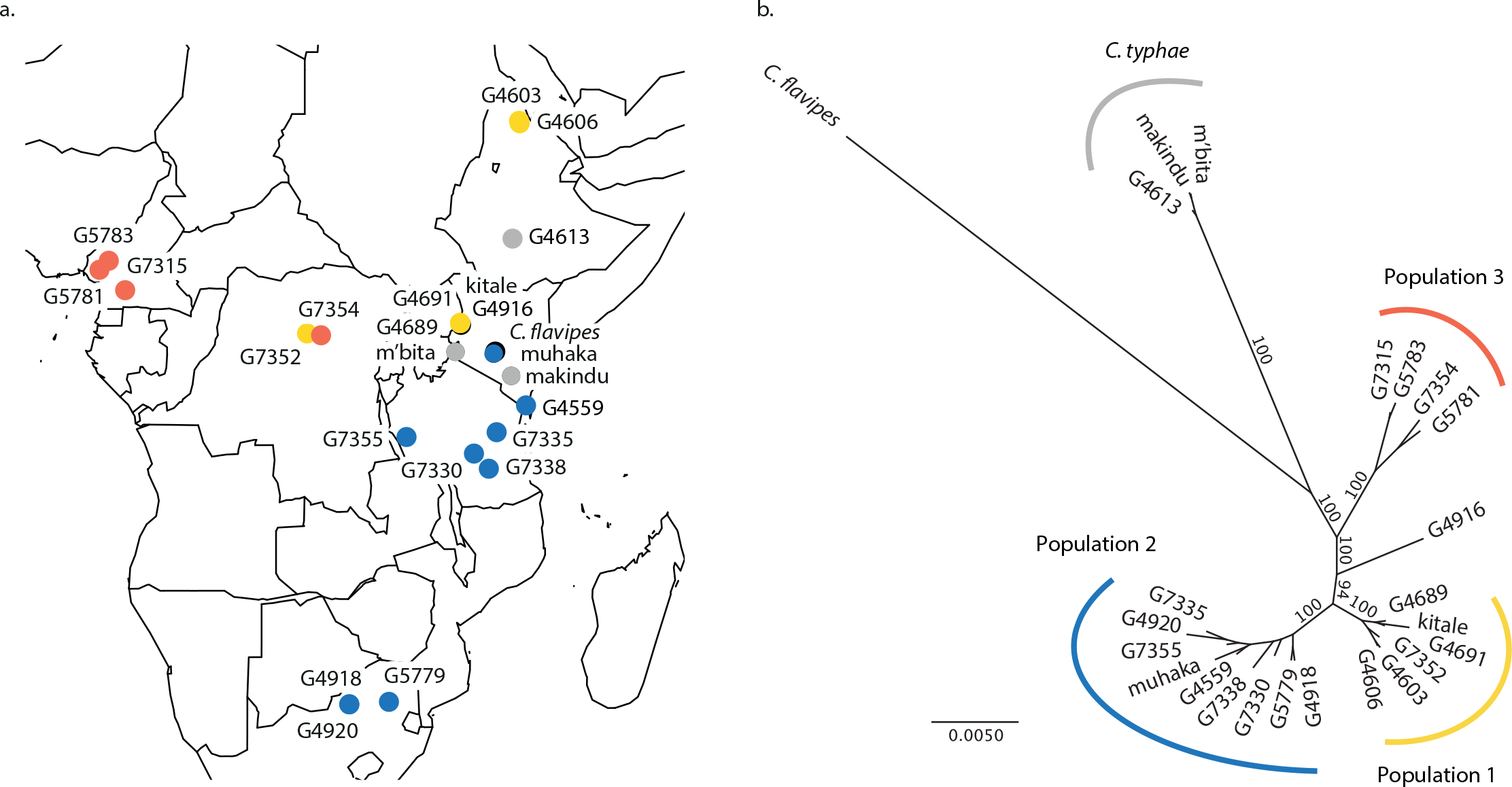
Geographic distribution and Phylogeny of *C. sesamiae* and *C. typhae* samples included in this study. a. Map of sampling sites, the colours represent the populations. b. Phylogenetic tree obtained with PhyML on all concatenated bracovirus genes and showing the relation between the samples. Populations have been defined based on phylogenetic clusters. Values on branch indicate the 1000 bootstrap node support.

### Detection of selection on bracovirus genes

To study the mechanisms involved in the divergence between *C. sesamiae* populations and with *C. typhae* between them and with the elated species, several tools, such as measures of F_ST_, d_XY_ and dN/dS ratio, were used to investigate selection signatures on the 98 bracovirus genes captured in experiment.

As a first approach, we measured the relative differentiation (F_ST_) on bracovirus genes for each pair of populations. Overall mean F_ST_ values were high in all comparisons, ranging from 0.53 to 0.86. However, we observed quite different profiles along the length of the bracovirus loci depending on the comparison (Figure 2). In the comparison between population 1, 2 and 3, the F_ST_ values of each bracovirus gene were heterogeneous although they averaged 0.53, because they varied from 0 to 1 throughout the proviral loci. The comparisons of populations 1, 2 and 3 with *C. typhae* showed completely different profiles. The F_ST_ values are largely higher with averages between 0.80 and 0.86 illustrating the strong differentiation of *C. typhae*. Notably, this high F_ST_ level was detected all over the bracovirus, along the macrolocus as well as segment 7, which is physically distant in the genome (Bézier *et al.* 2013). In all pairwise comparison, no outlier gene could be identified, because genes with high F_ST_ values were not separated from the whole distribution.

**Figure 2.**
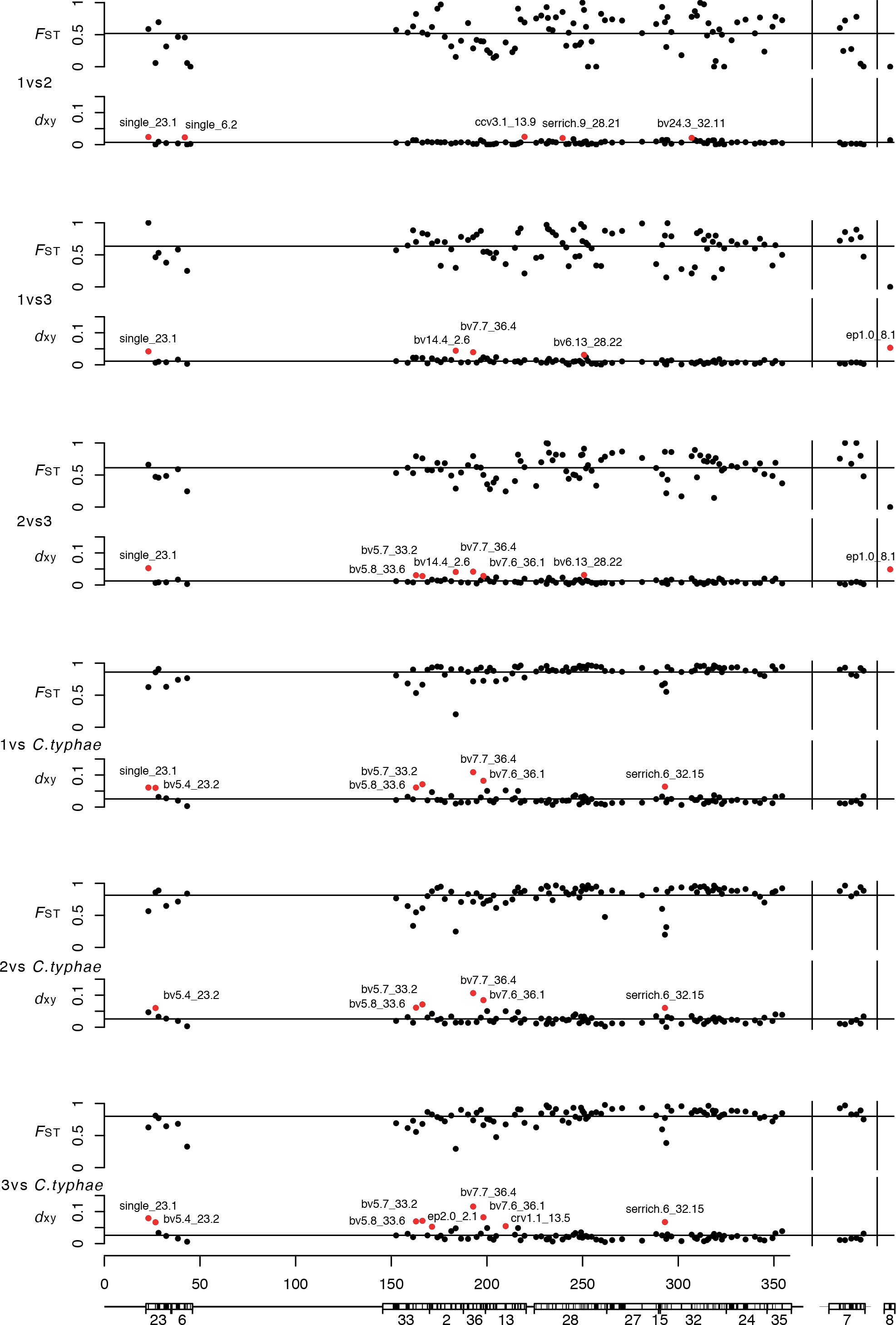
Differentiation of the bracovirus loci between different *C. sesamiae* populations and with *C. typhae*. Relative (F_ST_) and absolute (d_xy_) differentiation was measured for each gene. Horizontal lines indicate average levels. Genes with exceptional differentiation (outliers) are shown in red. The x-axis indicates the position in kb, note that there is a gap between 50 kb and 150 kb. Below, a graphical representation of the bracovirus with segments and genes in black highlighting their genomic organization.

As a second approach, the absolute differentiation (d_xy_) was measured between pairs of populations for each of the 98 bracovirus genes (Figure 2). The d_xy_ measures were quite different from the F_ST_. Mean d_xy_ levels were low in the comparisons between all the populations (ranging from 0.007 to 0.026) and were correlated with phylogenetic distances. In all comparisons some genes could be defined as outliers clearly separated from the general d_xy_ distribution. Out of the 98 bracovirus genes under focus, 16 genes, distributed throughout the bracovirus region, showed high genetic differentiation in pairwise population comparisons. Some genes were found as outliers in a single comparison. This is the case for *single_6.2, ccv3.1_13.9, serrich.9_28.1* and *bv24.3_32.1*, which were detected in the comparison between the closely related populations 1 and 2, as well as *ep2.0_2.1* and *crv1.1_13.5* found in the comparison between populations 3 and *C. typhae* (Figure 2). In these cases the d_xy_ alone is not able to determine if gene divergence results from adaptive divergence. In contrast some genes were found as outliers in all pairwise comparisons of particular populations, strongly suggesting that they might be implicated in the divergence of these populations. This is the case for five genes of *C. typhae* (*bv5.4_23.2, bv5.8_33.6, bv5.7_33.2, bv7.6_36.1 and serrich.6_32.15;* Figure 2). The *bv14.4_2.6, bv6.13_28.22* and *ep1.0_8.1* genes seemed to diverge specifically in population 3. Lastly, two genes, *single_23.1* and *bv7.7_36.4*, were defined as outliers in almost all comparisons. Interestingly, almost all the outlier genes belong to 6 out of 31 multigenic families of the bracovirus (families bv5, bv6, bv7, bv14, ep1 and serrich). Strikingly, within the targeted region, 3 out of 4 copies of the bv5 family and 2 out of 3 copies of the bv7 family were defined as outliers, suggesting both gene families might be particularly adaptive.

To determine if the outlier genes found in the d_xy_ analysis were undergoing diversifying selection, dN/dS ratio were estimated using the Branch-Site REL model implemented in HyPhy. Positive selection signatures were found in six out of sixteen outlier genes (Table 1). The *ep1.0_8.1* gene showed high dN/dS ratio in population 3. With five out of six genes evolving under positive selection, *C. typhae* lineage bore most of the hallmarks of molecular differentiation from *C. sesamiae*. Strikingly, two genes of the bv7 family (*bv7.7_36.4* and *bv7.6_36.1*) were found to be adaptive through this approach, as well as *bv5.8 33.6*, *ep2.0 2.1* and *crv1.1_13.5* (Table 1).

**Table 1:** Genes under positive selection in *Cotesia typhae* and *Cotesia sesamiae* populations. Genes are ordered by position in the genome; branch leading to the populations in which the gene is undergoing selection; value of the mean dN/dS ratio; significance estimated using LRT approach, p-value and p-value corrected using Holm-Bonferroni method for multiple comparisons.

| Gene | Branch | Mean dN/dS ratio | LRT | p-value | corrected p-value |
| --- | --- | --- | --- | --- | --- |
| ben.9_23.5 | 1 | 1.98 | 12.35 | 0.0007 | 0.0278 |
| ben.9_23.5 | 2 | 10.00 | 10.78 | 0.0016 | 0.0599 |
| single_6.2 | 2 | 0.93 | 7.59 | 0.0079 | 0.3323 |
| ben.11_33.5 | *C. typhae* | 1.50 | 8.93 | 0.0040 | 0.1725 |
| bv5.8_33.6 | *C. typhae* | 10.00 | 10.81 | 0.0015 | 0.0712 |
| bv5.6_33.1 | *C. typhae* | 2.79 | 8.14 | 0.0060 | 0.2761 |
| ep2.0_2.1 | *C. typhae* | 2.85 | 5.90 | 0.0187 | 0.8232 |
| single_2.3 | *C. typhae* | 10.00 | 6.56 | 0.0134 | 0.6146 |
| bv2.1_2.5 | *C. typhae* | 10.00 | 14.22 | 0.0003 | 0.0127 |
| bv7.7_36.4 | *C. typhae* | 2.30 | 12.19 | 0.0008 | 0.0356 |
| bv7.6_36.1 | *C. typhae* | 2.79 | 9.85 | 0.0025 | 0.1161 |
| ep2.2_13.1 | *C. typhae* | 7.58 | 6.98 | 0.0108 | 0.4964 |
| single_13.3 | *C. typhae* | 0.71 | 12.32 | 0.0007 | 0.0332 |
| bv8.3_28.14 | 2 | 1.09 | 7.47 | 0.0084 | 0.3707 |
| ben.13_27.2 | *C. typhae* | 8.94 | 9.12 | 0.0036 | 0.1637 |
| ben.13_27.2 | 3 | 2.90 | 11.38 | 0.0012 | 0.0534 |
| single_32.18 | *C. typhae* | 10.00 | 9.10 | 0.0037 | 0.1692 |
| ben.14_24.5 | *C. typhae* | 1.26 | 23.00 | 3.4e-06 | 0.0002 |
| crp.4_35.3 | *C. typhae* | 10.00 | 6.06 | 0.0173 | 0.7943 |
| ep1.0_8.1 | 3 | 2.76 | 17.47 | 5.4e-05 | 0.0046 |

### Pseudogenization and gene loss

The manual investigation of bracovirus gene alignments revealed fixed nonsense mutations as well as several deletions observed in all individuals from a given population. These mutations shortening the encoded proteins suggest on-going pseudogenization process. Surprisingly, pseudogenization events fixed in at least one population were found in at least 21 genes among the 98 genes studied. Nineteen genes harboured point mutations or deletion causing premature stop codons causing shortened ORFs. While some mutations reduced only a small part of the protein (e.g. bv5.8_33.6 lost 13% of the reference protein), others impacted considerably protein length (e.g. bv6_32.19 lost more than 85% of the protein; Table 2). Although pseudogenization events were found in all *C. sesamiae* populations, the specialist *C. typhae* lineage was most affected (Table 2).

**Table 2:** Bracovirus genes pseudogenized in particular populations. Genes are ordered by position in the genome; populations in which the mutations were fixed; type of mutation; size of wild type and shortened mutant ORFs

| Gene | Population | Mutations | Size of wild type ORF | Size of mutant ORF |
| --- | --- | --- | --- | --- |
| single_23.1 | *C. typhae* | 270bp Deletion | 183 | 93 |
| ben.3_6.3 | 3 | 7bp Deletion | 616 | 129 |
| single_6.2 | 3 | 255bp Deletion | 324 | 239 |
| single_6.2 | *C. typhae* | 619bp Deletion | 324 | 117 |
| single_6.4 | 3, *C. typhae* | Whole Gene Deletion | 148 | 0 |
| ben.11_33.5 | *C. typhae* | STOP codon | 1081 | 554 |
| bv1.2_33.3 | 3 | STOP codon | 165 | 79 |
| bv1.2_33.3 | 2 | 7bp Insertion / Frame-shift | 165 | 95 |
| bv5.8_33.6 | *C. typhae* | STOP codon | 119 | 93 |
| bv5.6_33.1 | *C. typhae* | Mutation splicing site | 142 | 69 |
| ep2.0_2.1 | *C. typhae* | STOP codon | 378 | 373 |
| bv11.3_2.7 | *C. typhae* | STOP codon | 321 | 315 |
| single_13.6 | 3 | 194bp Deletion | 166 | 101 |
| bv22.2_28.19 | 3 | STOP codon | 88 | 50 |
| serrich.9_28.21 | *C. typhae* | Deletion 2bp | 129 | 126 |
| bv8.4_28.11 | 2 | 1bp Deletion / Frame-shift | 133 | 48 |
| bv6.12_28.7 | *C. typhae* | STOP codon | 89 | 21 |
| bv6.13_28.22 | 3, *C. typhae* | STOP codon | 159 | 100 |
| bv23.1_32.19 | 1, *C. typhae* | STOP codon | 110 | 27 |
| bv23.1_32.19 | 2 | STOP codon | 110 | 16 |
| ep1.6_7.6 | 3, *C. typhae* | Whole Gene Deletion | 183 | 0 |
| ep1.0_8.1 | *C. typhae* | Whole Gene Deletion | 307 | 0 |

In a similar manner than for the locus 8 in *C. typhae*, the fine scale analysis of gene coverage revealed two large regions without any reads. To investigate if this absence of coverage along the bracovirus genome corresponded to real deletions, read mapping results were re-assembled to evaluate if both sides of these putative deletions were contiguous in these populations. In both cases, contiguous sequences bridging the deletion gap could be assembled with the same coverage as the rest of the genome, showing that these deletions were genuine. The deletions were found in the genes *single_6.4* and *ep1.6_7.6* (Figure 3). The deletion in the *single_6.4* gene was larger around the gene in the population 3 (1,056 bp deleted) and smaller in *C. typhae* (660 bp deleted). The situation was different in the *ep1.6_7.6* gene region with a smaller deletion in the middle of the gene for the population 3 (341 bp deleted) and a larger deletion including the whole gene for the *C. typhae* (608 bp deleted). As the positions of the deletions differed between the populations, they likely originated from independent events. In both cases the deletions were fixed, i.e. shared by all the samples, in populations 3 and *C. typhae*. Although the lack of flanking regions did not permit to conduct this analysis on the missing bracoviral locus 8 of *C. typhae*, these results suggest that there might also be a large deletion in that case.

**Figure 3.**
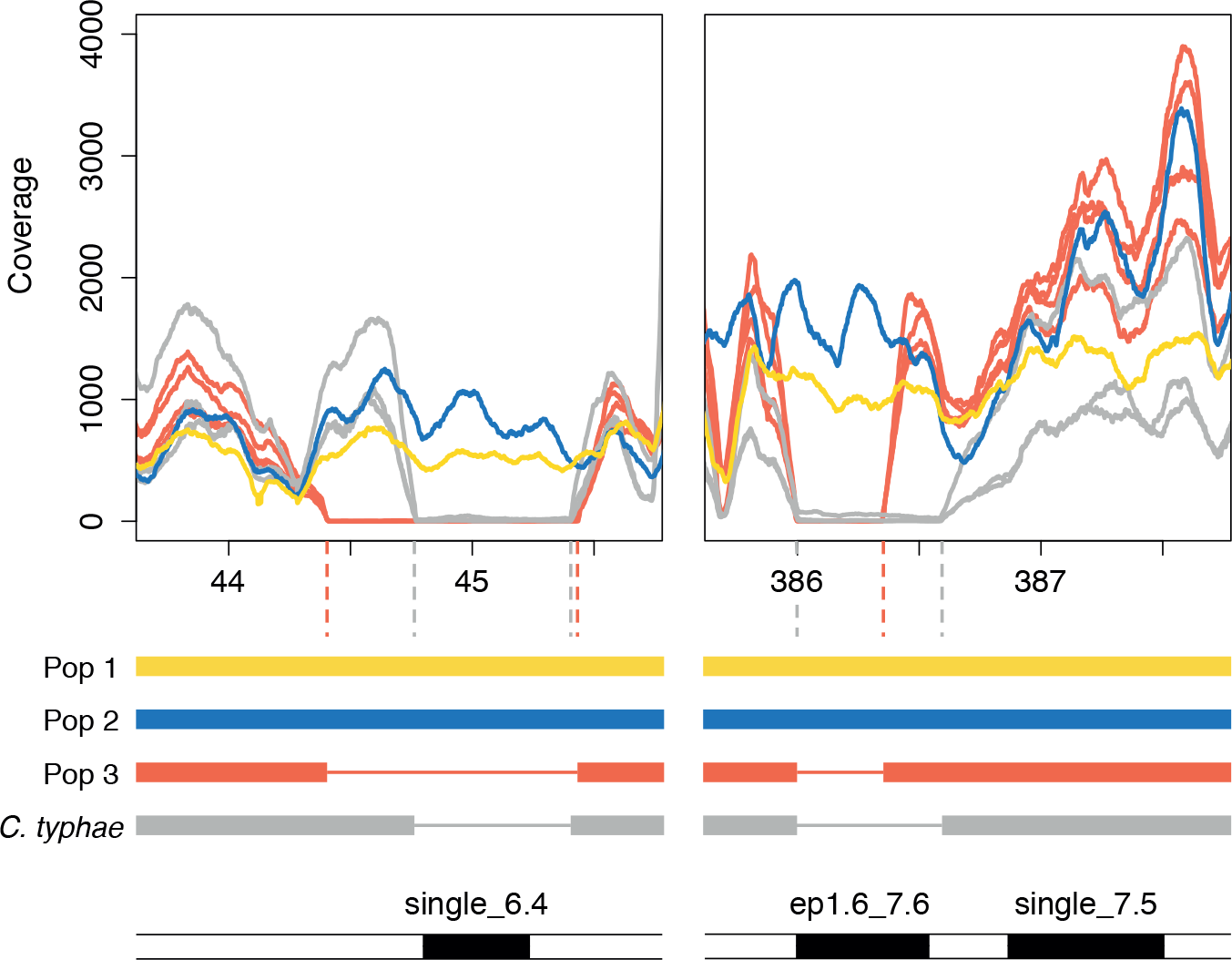
Read coverage plots of the whole deletions of the genes *single_6.4* and *ep1.6_7.6*. The coverage was reported for all samples showing the deletions and only one representative sample for the populations without the deletion. The x-axis indicates the position in kilo base pairs (kbp). Under the plots are represented illustrations of the genomic sequences for each population with a thin line on the deletion regions and in black the gene positions.

## Discussion

Generally used for the study of inter-specific genetic variability (Jones & Good 2016), the targeted sequencing approach has been particularly efficient to study the evolution of the bracovirus of *C. sesamiae* at the population level. The approach was validated through a series of tests evaluating its robustness (Chen *et al.* 2012), that showed the capture and sequencing were representative of the genetic diversity of the samples and populations studied. Furthermore the gene scale approach circumvented analytical bias resulting from the presence of numerous transposable elements bias in the bracovirus loci (Bézier *et al.* 2013).

### Distinct evolutionary lineages within the Cotesia flavipes species complex

The phylogenomic analysis based on all genes from the entire bracovirus showed a strong genetic structuration of the samples in five well-delimited clusters. The branch length and the differentiation level, F_ST_ and d_xy_, measured in the bracovirus region spanned different situations from closely related to deeply divergent clusters. This structuration is congruent to three clades identified in previous phylogenetic approach (Kaiser *et al.* 2015). The main difference is the basal position of the *C. typhae* species, which could be explained by the rapid evolution of the bracovirus genes leading to long branch attraction artefact in the phylogeny. A phylogeny based on another gene set might resolve these issue and clarify the phylogenetic position of *C typhae* within the *C. flavipes* species complex. Lastly, the same structuration in five clusters was found using a broad population genetic approach using neutral molecular markers on 638 samples from all over Sub-Saharan Africa (Branca *et al.* 2017) thereby confirming the reliability of the sample clustering based on 98 bracovirus genes.

*C. sesamiae* is found in Sub-Saharan Africa, and most populations occupy large geographic areas of cross-continental proportion (Branca *et al.* 2017). The existence of these large continuous populations requires to maintain genetic homogeneity through gene flow over long distances, and therefore large dispersion abilities. In Africa, *C. flavipes* species, belonging to the same species complex as *C. sesamiae*, has a dispersal rate estimated to range between 11 and 200 km per year (Assefa *et al.* 2008; Omwega *et al.* 2013). Similar dispersion abilities would contribute to the large population distribution observed in *C. sesamiae*. The populations appear broadly separated in Africa, with population 1 in the North-East, population 2 in the South and population 3 in the West. This structuration and the divergence observed could partly result from past geographic isolation. However population 1, 2 of *C sesamiae* and *C. typhae* do overlap in Kenya suggesting population divergence might be explained by other evolutionary forces, such as ecological adaptation.

### Adaptive evolution in the bracovirus loci

Host parasite relationship is the main evolutionary force that explains *C. sesamiae* population divergence and structuration (Branca *et al.* 2017). The ability to infest one host or another depends on the expression of particular virulence genes lifting particular host resistance (Elrich & Raven 1964). In this context the bracovirus, which functions as the transfer agent of several hundred virulence genes, e.g. 222 genes in *C. congregata* bracovirus (Bézier et al. 2013), plays a key role in the molecular interactions between parasitoid wasps and their hosts (Burke & Strand 2012; Drezen *et al.* 2014),. This should lead to ecological adaptation (Herniou *et al.* 2013; Gauthier *et al.* 2017). We therefore endeavoured to scan the bracovirus loci for genetic signatures of molecular adaptation between *Cotesia* populations and species with different host ranges.

As F_ST_ values for each bracovirus genes were extremely variable among population comparisons and particularly high in the comparisons with *C. typhae*, the identification of specific divergent genes within the bracovirus loci was not possible. There were however many genes with F_ST_ values above 0.8, suggesting undetected ongoing processes. This lack of results are in line with the recent criticisms on F_ST_ differentiation measures stating their non-independence to within-population diversity (Cruickshank & Hahn 2014). In contrast measures of the absolute divergence (d_xy_) between bracovirus genes in each population revealed low levels of overall genetic differentiation, as expected between populations, but highlighted several outlier genes, which were significantly more divergent. The level of differentiation between *C. typhae* and *C. sesamiae* populations remains relatively high, compared to compared sister species of *Heliconius* or *Anopheles* (Nadeau *et al.* 2012; Cruickshank & Hahn 2014). The differences observed between absolute and relative differentiation could result from the population history or the types of selection., Population divergence with or without gene flow or recurrent versus strong selection may indeed induce variation in allelic frequency and genetic diversity (Cruickshank & Hahn 2014; Irwin *et al.* 2016). To further discriminate between these evolutionary processes, information on allelic frequency all along the genome would be required. However working on virulence genes from a specific region allowed studying molecular evolution between *C. sesamiae* populations and *C. typhae* in finer details with dN/dS ratio comparisons to infer positive selection.

The no *a priori* approach combining different tools on 98 bracovirus genes highlighted six particularly divergent genes (*ep1.0_8.1*, *ep2.0_2.1*, *single_6.2*, *bv5.8_33.6*, *bv7.7_36.4*, *bv7.6_36.1*). Previous studies mainly focused on particular virulence genes, e.g. *ptp* (Serbielle *et al.* 2012), *Crv1* (Dupas *et al.* 2008) at the inter- and intra-specific level. The *Crv1* gene is involved in the inactivation of host haemocytes (Asgari & Schmidt 2002; Kumar & Kim 2014) and different genotypes were previously linked to to the abilities to infect *B. fusca* (Gitau *et al.* 2007; Dupas *et al.* 2008; Branca *et al.* 2011). However, when analysed conjointly with other bracovirus genes, it did not show any strong outlier signal. In our results, *Crv1* is divergent only in the comparison between the population 3 and *C. typhae* (Figure 3.) but does not show signature of positive selection (Table 1). This suggests previous host specific genotype observations might have resulted from hitchhiking rather than direct selection of the *Crv1* gene and that the overall host range of each population might lead to confounding molecular evolution results. Our results on *bv6*, *ep1* and *Serine-rich* (*serrich*) converge with a previous study of bracovirus genes at the inter-specific level including *C. congregata*, *C. vestalis* and *C. sesamiae* (Jancek *et al.* 2013). These genes showing signatures of positive selection, both in micro and macroevolutionary analyses, therefore appear important for the ecological adaptation of the *Cotesia* genus. This suggests the recurrent molecular evolution of the bracovirus driven by cycles of arms race between the parasites and the hosts (Pennacchio & Strand 2006). Furthermore, the endoparasitoid way of life could also play a role in the strength of the selection pressure imposed by the host on the parasitoids. These interactions are mainly mediated by bracoviruses, which are essential for larvae to overcome host immune defences and development, hence for the overall parasitic success of the wasp (Beckage *et al.* 1994).

The gene *CcV3.1*, diverging only between the populations 1 and 2, has been acquired by different lepidopteran species through horizontal transfer from the bracovirus. Functional analyses described its role in the caterpillar immune defenses against exogenous viral pathogens (Gasmi *et al.* 2015). Thus the function of this bracovirus gene may extend beyond the antagonistic relationship between host and parasite to interact within larger multitrophic frameworks including other pathogens.

In the candidate gene list, the *bv5* and *bv7* multigenic families are particularly well represented. These genes of unknown function are specific to *Cotesia* species (Burke & Strand 2012). In *C. congregata*, variation in gene expression of different *bv5* genes resulted from modifications in promoter region, due to transposable element insertion or genes presence (Chevignon *et al.* 2014). These variations in gene use within a gene family could have direct consequences on the biochemical interactions between parasite and hosts. Depending on the outcome and the ecological context in which the population live, different evolutionary pressures should be put on each gene within a family. Further functional analyses would be required to reveal how the six candidate genes interact with the different hosts and how they modulate host physiology leading to the wasp parasitic success. Moreover, in the context of biological control programs, these candidate genes, which are differentiated between populations, could be used as molecular markers, to identify the adapted wasp for an introduction against a targeted host and then to follow the acclimatization of these introduced wasps in the field.

### Evolutionary history of bracovirus genes

The bracovirus genes are organized in different multigenic families and certain genes in these families are under divergent selection. In several multigenic families, e.g. *bv5*, *bv7* or *ep1*, some genes show mutations inducing frame-shifts and stop codon reducing the protein size. These events impacting protein function result from relaxation of a negative selection and fixation of deleterious mutations. Large deletions also cause another type of less frequent gene loss. The *ep1* multigenic family, composed of three variants, follows these different evolutions. *Ep1.0_8.1* shows specific divergent selection pressure in the population 3, so this apparently adapted variant should be involved in successful parasitism of local hosts. In parallel, the *ep1.6_7.6* gene appears is completely deleted in population 3. This loss suggests it had become useless or was impeding parasitism success. This process of multigenic family evolution has been described by the ‘birth and death’ model (Nei & Rooney 2005). These gene families result from duplication events, including previously described large genomic duplication and rearrangements (Bézier *et al.* 2013), that creates new bracovirus genes, which become the supports for new mutations (Herniou *et al.* 2013; Francino 2005). If the mutations are adaptive, these versions will be submitted to various selection pressures that lead to different evolutionary trajectories as observed. In the case of specialization, only the most adapted versions are useful and the others experience a relaxation of the conservative selection pressures. This allows the accumulation of mutations that alter the function and lead to pseudogenization (Herniou *et al.* 2013, Gauthier *et al.* 2017).

### Host specialization

The use of well characterized natural populations, presenting contrasted ecological phenotypes (Dupas *et al.* 2008; Branca *et al.* 2011; Kaiser *et al.* 2015), gives a good framework to infer the evolutionary processes acting on these populations. *C. sesamiae* populations highlighted variations in host ranges, for example population 1 and 2 differ in the ability to infest *B. fusca* (Dupas *et al.* 2008; Branca *et al.* 2011). Moreover, *C. typhae* is strictly associated to one host *S. nonagrioides* (Noctuidae), all the samples (N= 46 in Branca *et al.* 2017 and N= 35 in Kaiser *et al.* 2015) have been collected on this species revealing strict specialization. This mechanism has been mainly described in phytophagous species (Via 2001; e.g. Groman & Pellmyr 2000; Peccoud *et al.* 2009) but is predicted to be as widespread in parasitoids (Rice 1987).

The adaptation to a specific host, when parasitoids have a higher fitness on this host species, is driven by selection pressures imposed by host on parasitoid population and boosted by the intimate relationships that exist between the parasitoid and its hosts. The signatures of these selection pressures impact genes involved in host/parasite interaction and correspond to the signals identified in the virulence genes (Thompson 2005). Under certain condition this specialization process induce the ecological speciation and the emergence of a new species. Whole genome analyses could provide some keys to elucidate the speciation process involved in the emergence of *C. typhae*. The ecological speciation process, in sympatry, induce particular patterns located in specific genomic loci called ‘genomic islands’, such as described in stick insects (Nosil *et al.* 2009; Yeaman & Whitlock 2011; Nosil & Feder 2011; Feder *et al.* 2012). The high level of differentiation observed along the bracovirus suggests it might evolve as such a genomic island of divergence. The comparisons between the bracovirus and the rest of the genome would settle the hypothesis. Moreover, other adaptive traits could be involved in the evolution of these populations. For example, *C. typhae* is also specialized on one main plant *T. domingensis* (Kaiser *et al.* 2015). The wasp ability to detect host and host plant could also be involved in the specialization mechanisms. Different studies highlight the intimate relationship between plant and *Cotesia* parasitoids, which are mediated by plant volatiles induced by herbivore attacks (Geervliet *et al.* 1994). A genome scan approach to evaluate the molecular signature of selection on the whole genome could identify other genes and functions potentially involved in the evolution of parasitoid wasps.

## Acknowledgments

We thank Corentin Paillusson and Antoine Branca for sharing preliminary population genetics analyses. This project was funded by CEA Genoscope (National Centre of Sequencing), the European Research Council grant GENOVIR and the ANR Bioadapt ABC-Papogen. We are grateful to the Genotoul bioinformatics platform for computing and storage resources.

